# Engineering ssRNA tile filaments for (dis)assembly and membrane binding

**DOI:** 10.1101/2022.10.18.512742

**Authors:** Nicola De Franceschi, Baukje Hoogenberg, Cees Dekker

## Abstract

Cytoskeletal protein filaments such as actin and microtubules confer mechanical support to cells and facilitate many cellular functions such as motility and division. Recent years have witnessed the development of a variety of molecular scaffolds that mimic such cytoskeletal filaments. Indeed, filaments that are programmable and compatible with biological systems may prove useful in studying or substituting such proteins. Here, we explore the use of ssRNA tiles to build and modify cytoskeletal-like filaments *in vitro*. We engineer a number of functionalities that are crucial to the function of natural cytoskeletal systems into the ssRNA tiles, including the abilities to assemble or disassemble filaments, to tune the filament stiffness, to induce membrane binding, and to attach binding proteins. The work paves the way for building dynamic cell-like cytoskeletal systems made out of rationally designed ssRNA tiles that can be transcribed in natural or synthetic cells.

## Introduction

Cells feature a variety of cytoskeletal proteins that arrange into filamentous structures that are responsible for conferring mechanical properties to the cell. These systems are key to sustaining cellular functions such as cell shape, cell motility, intracellular transport, and cell division. Examples include actin^1^, tubulin^2^, ESCRT-III^3^, and the bacterial FtsZ^4^ and archaeal Cdv^5^. All of these form filaments that act as recruiting hubs for many other proteins that work in concert with the membrane. A variety of molecular scaffolds have been developed that mimic such cytoskeletal filaments, for example for *in vitro* studies with reconstituted minimal systems. Notably, in bottom-up synthetic biology, people have recently started efforts to build synthetic cells with various cellular components including natural cytoskeletal filaments and mimics thereof^6^. In such an approach, the use of natural proteins presents obvious advantages to create a cytoskeleton but it also has drawbacks such as a significant burden on the protein translation system as well as challenges in fine-tuning protein structure and function and encoding new functionalities. It is therefore of interest to study complementary systems that mimic cytoskeletal filaments which may provide advantages as substitutes for mechanical stability and cell division of future synthetic cells. The development of rationally designed tuneable synthetic scaffolding systems is of great interest, in particular if these can be functionalized similar to biological cytoskeletal systems.

Nucleic acids nanotechnology allows the rational design of nanoscale objects that are able to self-assemble into programmable shapes ranging from smileys^7^ to transmembrane channels^8,9,10^ and rotary motors^11,12^. In mimicking cytoskeletal systems, a natural approach is to assemble synthetic filaments made of tiles. Such tiles can be composed either by multiple strands of ssDNA^13^ or ssRNA^14^. Advantages of this approach include the presence of multiple 3’ and 5’ free ends, which allows to easily functionalize the tiles. However, such a design requires both the maintenance of a precise stoichiometry of all the strands composing the tiles and their folding by thermal annealing, which may limit their range of applications. A different approach is based on self-folded ssRNA tiles that are formed from a single ssRNA molecule and that can interact with each other via kissing loops (KLs)^15^. This presents a number of advantages such as the lack of restraints in terms of stoichiometry and the ability to fold co-transcriptionally at room temperature. Such ssRNA scaffolds have great potential to be expanded for a variety of applications, as can for example be modified to encode a range of curvatures^16^ and can be scaled up to yield kilobase-sized filaments^17^. Moreover, ssRNA molecules can be directly transcribed from the genome and thus are fully cell biology compatible. So far however, only limited attempts have been pursued to expand their capabilities by functionalizing the ssRNA scaffolds, e.g. by going beyond static structures by engineering dynamic filaments that can assemble as well as disassemble in a controlled way.

Here, we assemble and functionalize ssRNA tile filaments, and we demonstrate that we are able to tune their stiffness, control their polymerization, encode direct membrane binding, and create binding sites for proteins. We introduce additional designing principles, e.g., we design softer tiles that allow filament disassembly by strand displacement. Our work provides a toolbox that paves the way for using ssRNA tile filaments in dynamic cell-like systems.

## Results and discussion

Our starting point was previous work by on ssRNA tiles^15^. We made a 2-helix ssRNA origami tile design with “Antiparallel Even” (AE) crossovers by substituting the 120ᴼ Kissing Loops (KLs) present at the extremities of each helix with 180ᴼ KLs, so that tiles can polymerize in a head-to-tail fashion. This initial tile design is hereafter named T1 and it is detailed in Figure 1A, whereas we will also introduce other tiles variants with different modifications and functionalities. Assuming dsRNA is in A form^18^, the designed tile dimensions were 18.2 nm x 4.8 nm x 2.4nm (length x width x height). Tiles were folded by thermal annealing, deposited on mica and imaged by Atomic Force Microscopy (AFM).

**Figure 1:**
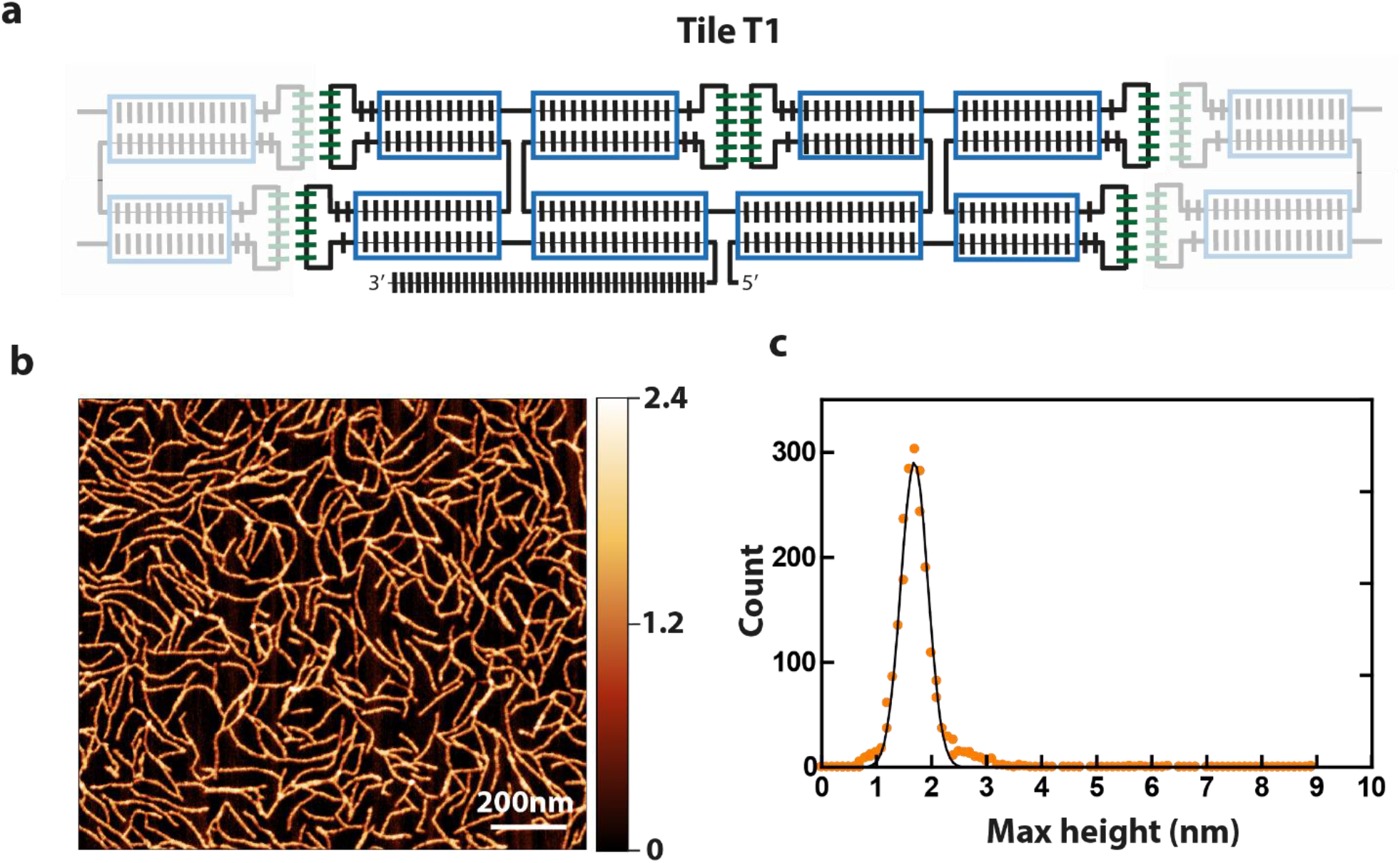
Design of ssRNA tile filaments. **(a)** Design of a T1 tile. Each bar corresponds to one base. Green bars indicate the 180ᴼ KLs. Bars outside the blue rectangles indicate unpaired bases. The shaded tiles on the sides illustrate how tile-tile interaction is achieved by pairing of the KLs. **(b)** AFM scan of T1 tiles that self-assembled into long linear filaments upon thermal annealing. **(c)** Quantification of filament height by AFM line scan analysis. A Gaussian fit is indicated by the black line.

The KL title-tile interactions resulted in the formation of long linear filaments, see Figure 1B. These filaments featured a submicron length and the height was 1.7 nm ± 0.3 nm, as measured by using dry AFM which provides a lower limit (Figure 1C). Residual monomeric tiles were not observed, indicating that the tile-tile interaction mediated by merely two KLs was very efficient, sequestering single tiles away into the filaments.

In order to visualize individual tiles within the filaments, we modified the design by introducing a 3-way-junction motif from the bacteriophage Phi29 DNA packaging motor^19^, which forms a rigid branching at a ≈60ᴼ angle within the helix that it originates from^20^. These tiles, named T2 (Figure 2A) can be identified within the filament due to the distinctive spike created by the 3-way junction (Figure 2B). This allowed to quantify the number of tiles present in each filament and estimate an average tile length of 18.8 ± 1.5 nm (mean ± SD; n=54), which is in very good agreement with the expected length of 18.2 nm. T2 filaments exhibited an average length of 148 ± 6 nm (mean ± SEM; n=623) and a persistence length of 46 ± 2 nm (mean ± SEM; n=623) (Figure 2E, 2F). Notably, the persistence length is close to that of dsDNA (∼50 nm^21^).

**Figure 2:**
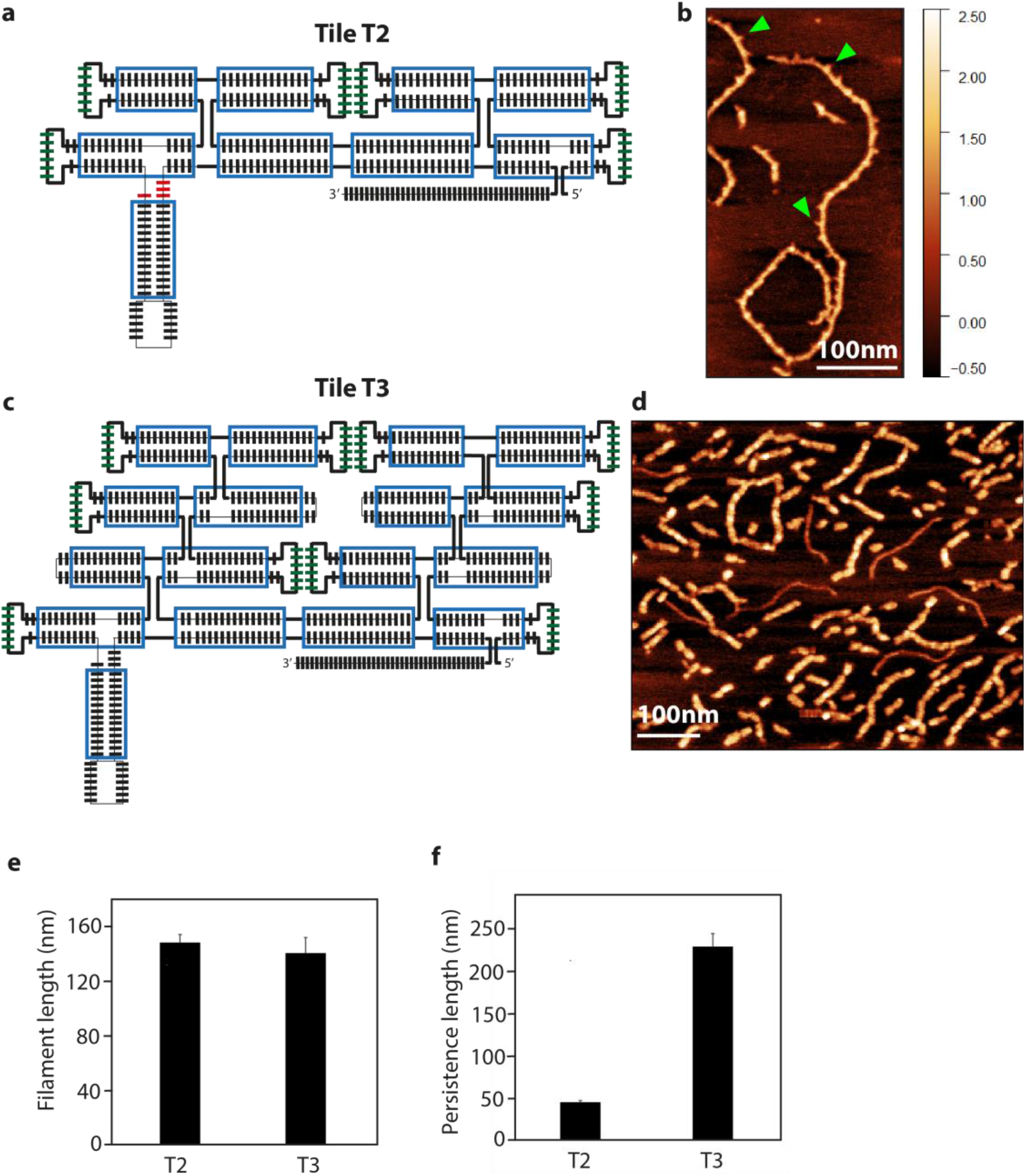
Tuning the persistence length of ssRNA tile filaments. **(a):** Design of tile T2. The bases forming the 3-way-junction motif are indicates in red. **(b)** AFM image of filaments made of tiles T2. Green arrowheads indicate spikes produced by the extra helix. **(c)** Design of tile T3. **(d)** Filaments of tile T3 imaged by AFM. Some additional thinner filaments are visible which are the ssRNA of unfolded tiles. **(e)** Quantification of length of T2 and T3 tile filaments. **(f):** Quantification of the persistence length of T2 and T3 tile filaments.

We sought to increase filament stiffness by modifying the design with the addition of 2 helices (named T3, Figure 2C). The resulting larger tiles formed filaments (Figure 2D) with a similar length as the T2 design (141 ± 11 nm (mean ± SEM; n = 149), but with a markedly higher persistence length of 229 ± 15 nm (mean ± SEM; n=149; Figure 2E, 2F). The design thus successfully allowed to tune filament stiffness to a 5-fold higher value.

Next, we engineered a membrane-binding interaction to the tiles to induce attachment of the filaments to a bilayer lipid membrane. A common approach to bind nucleic acid nanostructures to membranes is the use of a short DNA oligomer that hybridizes to the DNA/RNA nanostructure and that is chemically functionalized with cholesterol moieties that anchors to the membrane^22^. We initially used this strategy to bind our filaments to the membrane, by indeed hybridizing a cholesterol-functionalized oligo to the 5’ end of each RNA tile. However, while this was partially successful, the filaments appeared to be only weakly attached to the membrane, as multiple passages of the AFM tip in imaging drastically reduced their number (Figure S1a). We found that we could significantly improve the stability of filaments-membrane binding by engineering a direct interaction between the tile and the lipid membrane. To this aim, we inserted biotin aptamers^23^ into the T3 tiles, yielding the T4 design (Figure 3A). These 3’-pCp-Cy5-labelled T4 tile filaments exhibited very robust binding to membranes (Figure S1b). Thus, they could be imaged extensively both by liquid AFM as well as with confocal microscopy (Figure 3B, 3C). Figure 3C clearly show an abundance of filaments on the lipid bilayer, whereas no filaments were seen to attach to the bare mica. In addition to simplifying the system, insertion of the biotin aptamer allowed a tight and close contact between filaments and the membrane, which mimics cytoskeletal proteins such as the ESCRT-III complex^24^.

**Figure 3:**
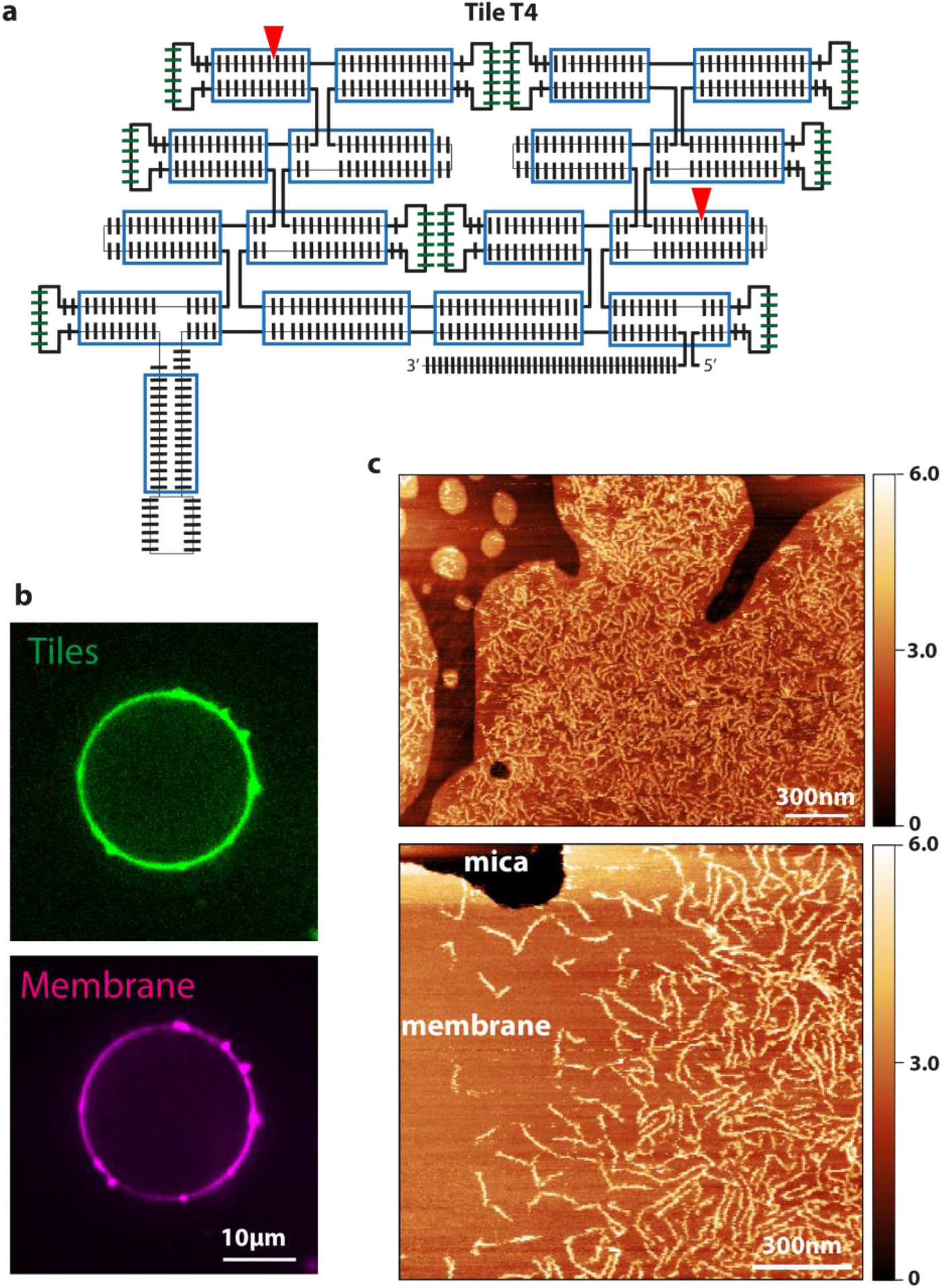
Engineering ssRNA tile filaments for membrane binding. **(a):** Tile T4 design. Red arrowheads indicate the positions where two biotin aptamers were inserted. **(b)** Tiles T4 binding to a giant liposome containing biotinylated lipids. The ssRNA tiles are indicated in green, the lipids in magenta. Single confocal plane. **(c)** Tiles T4 binding to lipid bilayer patches containing biotinylated lipids, as imaged by liquid AFM.

A common feature of cytoskeletal systems is their ability to be regulated by other proteins, which can either promote or inhibit filament polymerization^25^. In order to mimic such a functionality, we designed “capping tiles” (hereafter referred to as (T-cap”) that feature only one pair of KLs (Figure 4A). T-caps should effectively inhibit further polymerization of tiles T2. Indeed, we found that addition of T-caps completely abolished filament formation (Figure 4B). Another common feature of cytoskeletal filamentous systems is the presence of unstructured sequences that act as recruiting hubs for additional proteins that may modify the filament. An example is the linear C-terminal sequence of ESCRT proteins, which recruits the AAA ATPases Vps4 and CdvC^24,26^ which ultimately leads to filament disassembly. In our tile design, the 3’ extension represents a convenient site for recruiting enzymes such as helicases, that may be able to act on the filaments. Indeed, we observed a robust recruitment of the thermophilic helicase Hel308 to T2 filaments, see Figure 4C.

**Figure 4:**
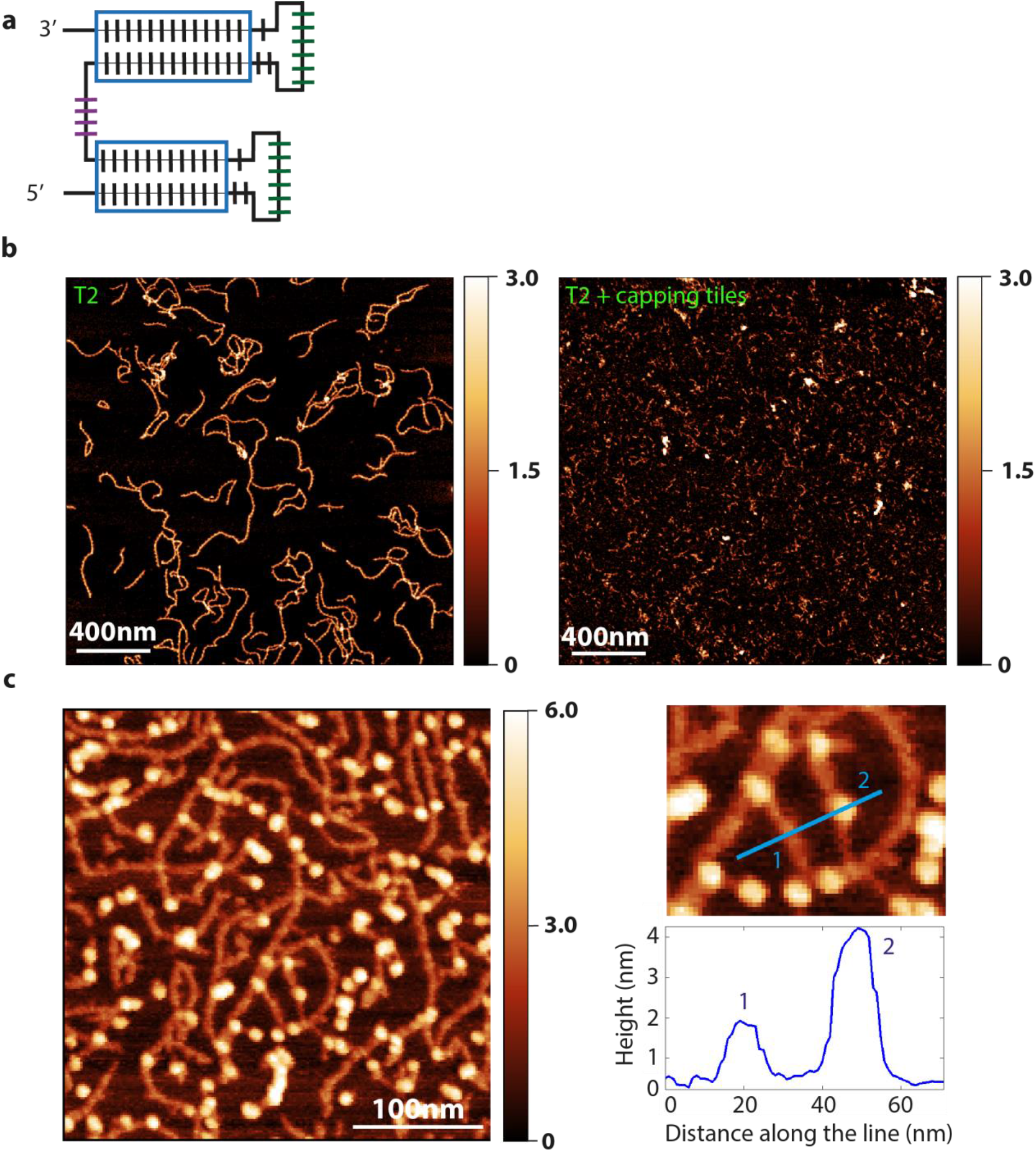
Inhibition of tile polymerization and engineering of a filament-protein interaction. **(a)** Design of T-cap. **(b)** Inhibition of T2 tiles polymerization by the addition of T-caps. **(c)** Helicase Hel308 binding to tile filaments. White blobs correspond to individual helicase molecules bound to the filaments. The right panel shows a magnification; the height profile across the blue line is shown in the bottom panel.

Finally, a crucial aspect of cytoskeletal filaments is their ability to undergo cycles of assembly and disassembly. While tile-tile interactions spontaneously drive filaments towards assembly in our system, we sought to engineer ways to disassemble them as well. Strand displacement is a well-established approach to disassemble DNA and RNA nanostructures^27^. In our design, the 3’ end can act as toehold, and an equimolar amount of reverse strand should be sufficient to disassemble filaments. However, we did not observe T1 filament disassembly upon addition of the T1 reverse strand, even after prolonged incubation (Figure 5A). Since this may be due to the high GC content of the tiles which leads to a very stable structure, we modified the original design in order to “soften” the tiles. Regions with high GC content (70-83%, indicated in red in Figure 5B) were lowered to a GC content of only 26-50%. Moreover, we added some unpaired bases between RNA helices (purple circles in Figure 5B). The resulting tiles (T5) were still able to form filaments (Figure 5C), but now the addition of the reverse T5 strand caused complete filament disassembly (Figure 5D) at room temperature.

**Figure 5:**
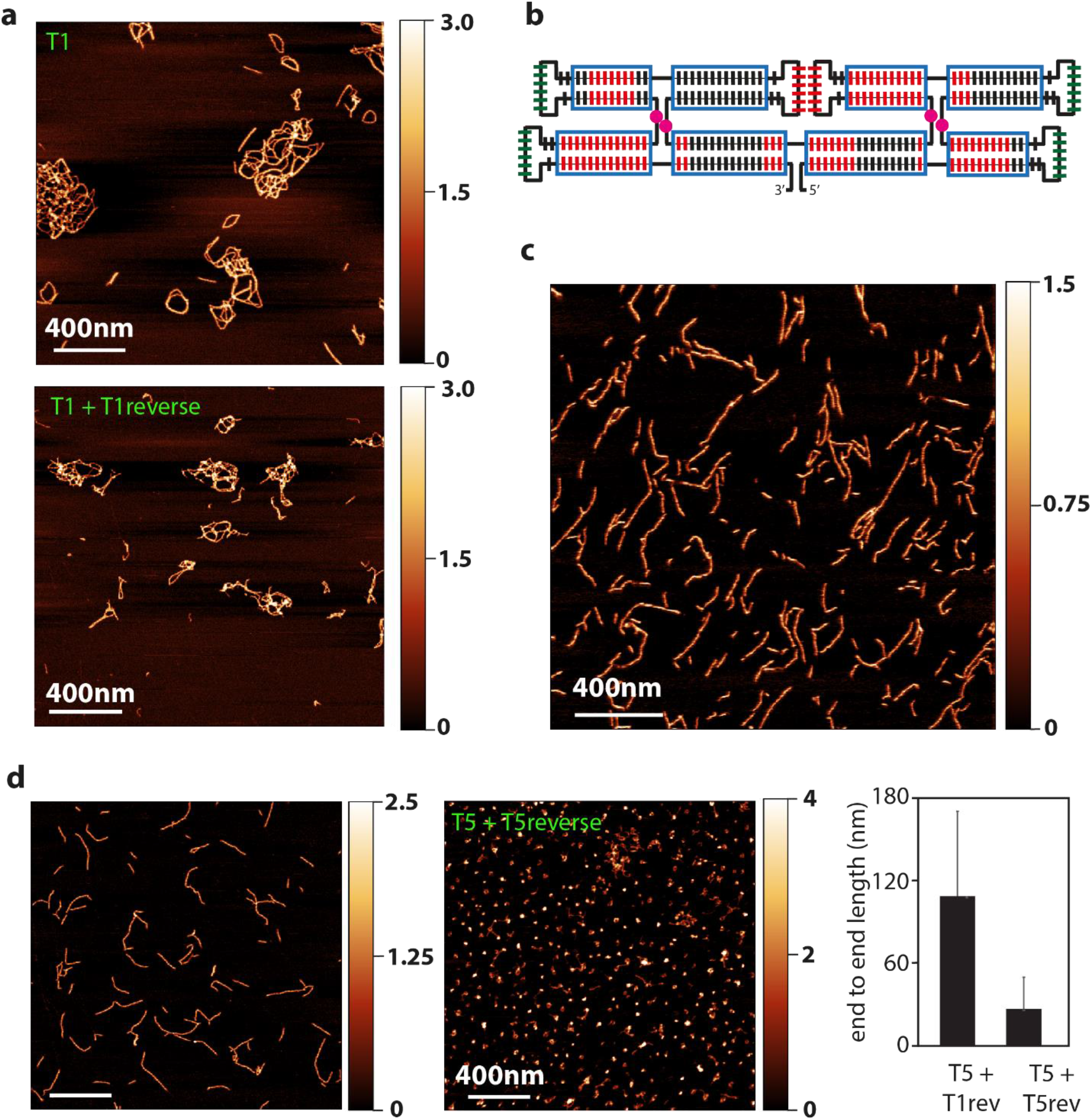
Engineering of filaments disassembly. **(a)** Filaments formed by tiles T1 (top) did not disassemble upon addition of equimolar amount of the T1 reverse strand (bottom). **(b)** Design of “soft” tiles T5. Red bars indicate regions of high GC content that were substituted by AT. Pink circles indicate extra unpaired bases that were added in between the helices. **(c)** AFM images of filaments formed by tiles T5. **(d)** Disassembly of filaments formed by tiles T5 upon addition of equimolar amount of T5 reverse strands.

## Conclusions

In this paper, we developed a range of functionalities on ssRNA tiles that facilitates the formation of cytoskeletal-like filaments. By changing the number of helices, we obtained filaments of variable stiffness. By branching the RNA sequence with specific motifs, we could engineer membrane binding via biotin aptamers, resulting in strong and direct binding. Moreover, we showed that binding sites for RNA-binding proteins such as helicases can easily be added with minimal modification of the design. Lastly, we modified the tile sequence to render them softer to achieve filament disassembly by strand displacement.

The use of self-folded ssRNA tiles presents a number of advantages over proteins, DNA, or multi-strand RNA tiles. An obvious advantage of self-folding ssRNA tiles is the possibility of directly producing them inside natural or synthetic cells by transcription, thus avoiding the need for a protein translation system. ssRNA tiles are furthermore able to undergo co-transcriptional folding at room temperature^15^. In our experiments we found that ssRNA tile can also undergo fast spontaneous isothermal refolding without the need to be coupled with transcription (Figure S2). Therefore, if a suitable enzyme-driven pathway for tile disassembly could be implemented, filaments made of tiles could in principle undergo cycles of assembly and disassembly driven by ATP consumption, rather than disassembly driven by strand displacement. Enzymes such as helicases, which we showed to be able to bind ssRNA tiles, and our improved “softer” design that facilitates disassembly, appear to be promising candidates that set the stage to build such system in future research.

In this paper, we showed how the ssRNA tiles design can be significantly expanded with a number of functionalizations, allowing parameters to be fine-tuned and providing a versatile platform for mimicking cytoskeletal filaments. Reconstituting the functionalities of cytoskeletal filaments is an important goal in bottom-up synthetic biology, where one of the most ambitious aims is the reconstitution of cell division^28^. The system we describe here paves the way for the development of molecular scaffolds that are programmable, dynamic and compatible with biological systems, on the route to the development of artificial cells.

## Acknowledgments

We thank Chenxiang Lin (Yale University) for discussions and training of N.D.F. in RNA design and handling, and Allard Katan and Alejandro Martin Gonzalez for assistance in AFM imaging. We acknowledge funding support from the BaSyC program of NWO-OCW and from the ERC Advanced Grant 883684.

## Methods

### Tiles sequence

The sequence of the different tiles designs are indicated as the coding strand on the artificial gene. The promoter region in underlined:

>T1

<u>TAATACGACTCACTATAGGG</u>AAGGAAGTGAGTAGTAGTCCACTGAGGGTGAAGAGCCTACGCCCTCGGTGGCGA

AGCGACCTGAAGCTCGCACGGGTCGTTTCGCGCAGGTGGCTGAAGCCTCCACGGCCATCTGCCACTGCTACTCGCTTCCGCGAAATGTCAATACGGACAGCGACGGTGAAGGAGGCACGCCGTTGCTGTCGACGGAGGCTGAAGCGAGCACGGCCTCTGTCGGGAGCTCTGCTGAAAGGCTCACGGCAGGGCTCCCCGTATTGGCATTTCGCAAAAACGTAGCATGCACAAAAGTAGCATCGACCCAGAGCAC

>T2

<u>TAATACGACTCACTATAGGG</u>AAGGGUGAAGAGCCUACGCCAUCGGUGGCGAAGCGACCUGAAGCUCGCACGG

GUCGUUUCGCGCAGGUGGCUGAAGCCUCCACGGCCAUCUGCGACUGCUACUCGCUUCCGCGAAAUGUCAAUACGGACAGCGACGGUGAAGGAGGCACGCCGUUGCUGUCGACGGAGGCUGAAGCGAGCACGGCCUCUGUCGGAUCAUCUGCUGAAAGGCUCACGGCAGGUGUGUAUGGUGAAGGUCAAAGCUCACAGAAAGACCUUCACCAUACUUUGUCCCGUAUUGGCAUUUCGCGGAAGUGAGUAGUAGUCCACUGAUGACAAACACACAAACACAAACACACAAACACCUGUGAGC

>T3

<u>TAATACGACTCACTATAGGG</u>AAGGTAGTCAGAACTTGTCGACGGACGCTGGTAACGGCGTCTGTCGGGTAGCCAC

GTGGCATCGCGTGGTTACTCCACTGAGGGTGAACCACGTACGCCCTCGGTGGCGAAGCGACCTGAACGTCCAACGGGTCGTTTCGCGCAGGTGGCTGAAGCCTCCACGGCCATCTGCGACTGCTACTCGCTAGAGAGTGAGTAGTAGCCG

GACGAGCCTGAAGCGTACACGGGCTTGTCCCACAGGTTCTGGCTACCCCGATATCTCTACACGGTGAGGGTCGTTGAAGTACGCACGACGATCCTCCCCGTATTGGCATTGTAAAATGTCAATACGGACAGCGACGGTGAAGGAGGCACGCCGTTGCTGTCGACGGAGGCTGAAGCGAGCACGGCCTCTGTCGGGAGCTCTGCTGAAAGGCTCACGGCAGGGCTCACGTCGCACGCGGTACGCTGCGTGTGACCGGTGGTCAGGTGAATGCGCCTGGCCACCGCGTGTAGGGATATCGG

>T4

<u>TAATACGACTCACTATAGGG</u>CUGAACGUCCAACGGCGUCUGUCGGGUAGCCACGUGGCAUCGCGUAAGGUCA

GGAACAAAACCGACCAUAGGCUCGGGUAUCGAAAAGCCUAUGAACAAACCUGACCAAGGUUACUCCACUGAGGGUGAAGAGCCUACGCCCUCGGUGGCGAAGCGACCUGAAGCUCGCACGGGUCGUUUCGCGCAGGUGGCUGAAGCCUCCACGGCCAUCUGCGACUGCUACUCGCUAGAGAGUGAGUAGUAGCCGGACGAGCCUGAAGCGUACACGGGCUUGUCCGACAGGUUCUGGCUACCCCGAUAUCUCUACACGGUGAGGGUCGUUGAAGUACGCACGACGAUCCUCCCCGUAUUGGCAUUGUAAAAUGUCAAUACGGACAGCGACGGUGAAGGAGGCACGCCGUUGCUGUCGACGAAGCUCACCACACAAACCGAUCAUAGGCUCGGGAACCGAAAAGCCUAUGACACAAGGUGAGCAAGAGGCUGAAGCGAGCACGGCCUCUGUCGGGAGCUCUGCUGAAAGGCUCACGGCAGGGCUCACGUCGCACGCGGUACGCUGCGUGUGACCGAUCAUCAGGUGAAUGGACGACGCCUGGUGUGUAUGGUGAAGGUCAAAGCUCACAGAAAGACCUUCACCAUACUUUGUCGCGUGUAGGGAUAUCGGGGUAGUCAGAACUUGUCGACGGACGACAAACACACAAACACAAACACACAAACACCUGUGAGC

>T5

<u>TAATACGACTCACTATAGGG</u>TATGAAAGAATGATACTCACTTCAGTGAAATGTCAATACTAAGACAGTGACAGTGA

AGTAGTCACACTGTTACTGTCTACAGAAGTTGAAGTGATCACAACTTCTGTAAAGTGATCTTTGCTAAAAGATTCATGGTAGAGATTACAGTATTGACATTTCATTGAGGTGAGTATTATTCCAATTAGATAGAAGAATCTACTATCTAGTTG

AAGAGAAGCTACTTGAAGATCACACGAGTAGTTTCTCTCATGTAGATGAAGACTACACGTCTAACAAACACACAACCACAAACACCCAAACAC

>T5reverse

<u>TAATACGACTCACTATAGGG</u>GTGTTTGGGTGTTTGTGGTTGTGTGTTTGTTAGACGTGTAGTCTTCATCTACATGAG

AGAAACTACTCGTGTGATCTTCAAGTAGCTTCTCTTCAACTAGATAGTAGATTCTTCTATCTAATTGGAATAATACTCACCTCAATGAAATGTCAATACTGTAATCTCTACCATGAATCTTTTAGCAAAGATCACTTTACAGAAGTTGTGATCACTTCAACTTCTGTAGACAGTAACAGTGTGACTACTTCACTGTCACTGTCTTAGTATTGACATTTCACTGAAGTGAGTATCATTCTTTCATA

>T-cap

<u>TAATACGACTCACTATAGGG</u>ACACTGAGGGTGAAGAGCCTACGCCCTCGGTGAAAAGCGAAGCGACCTGAAGCTC

GCACGGGTCGTTTCGC

### Tiles Folding

Tiles were folded in 40mM Tris-Acetate (pH 8.0) + 2mM EDTA + 12.5mM Mg-Acetate using the following thermal cycle: 5’ at 80ᴼC; from 80ᴼC to 70ᴼC, -1ᴼC/min, 10 cycles; from 70ᴼC to 22ᴼC, -0.2ᴼC/min, 240 cycles.

### Quantification of filaments

The height of the filaments was obtained by applying a threshold mask to the filament image, selecting all contiguous regions of height larger than 0.3 nm above the background and larger than 9 pixels (215 nm^2^). These regions consist of partial, complete or multiple filaments. The distribution of maximum heights of these regions was fitted with a normal distribution to obtain mean and standard deviation. Quantification of filaments length and persistence length was done in a semi-automatic way by using DNA trace version 3.4.

### AFM imaging

Images were taken with a multimode-2 AFM from Bruker (Bruker corporation, Germany) using Scanasist-Air-HR tips from Bruker. The AFM was operated using Peak-Force tapping mode for imaging in air, at room conditions. Image data and processing (plane subtraction and flattening) was done with Gwyddion software.

## Notes

### Competing Interest Statement

The authors have declared no competing interest.

