## Supplementary figures 1 and 2 for "Engineering ssRNA tile filaments for (dis)assembly and membrane binding"

### Table of contents

**Supplementary Figure 1:** tile binding to membrane via cholesterol or aptamer anchors

**Supplementary Figure 2:** isothermal tile refolding

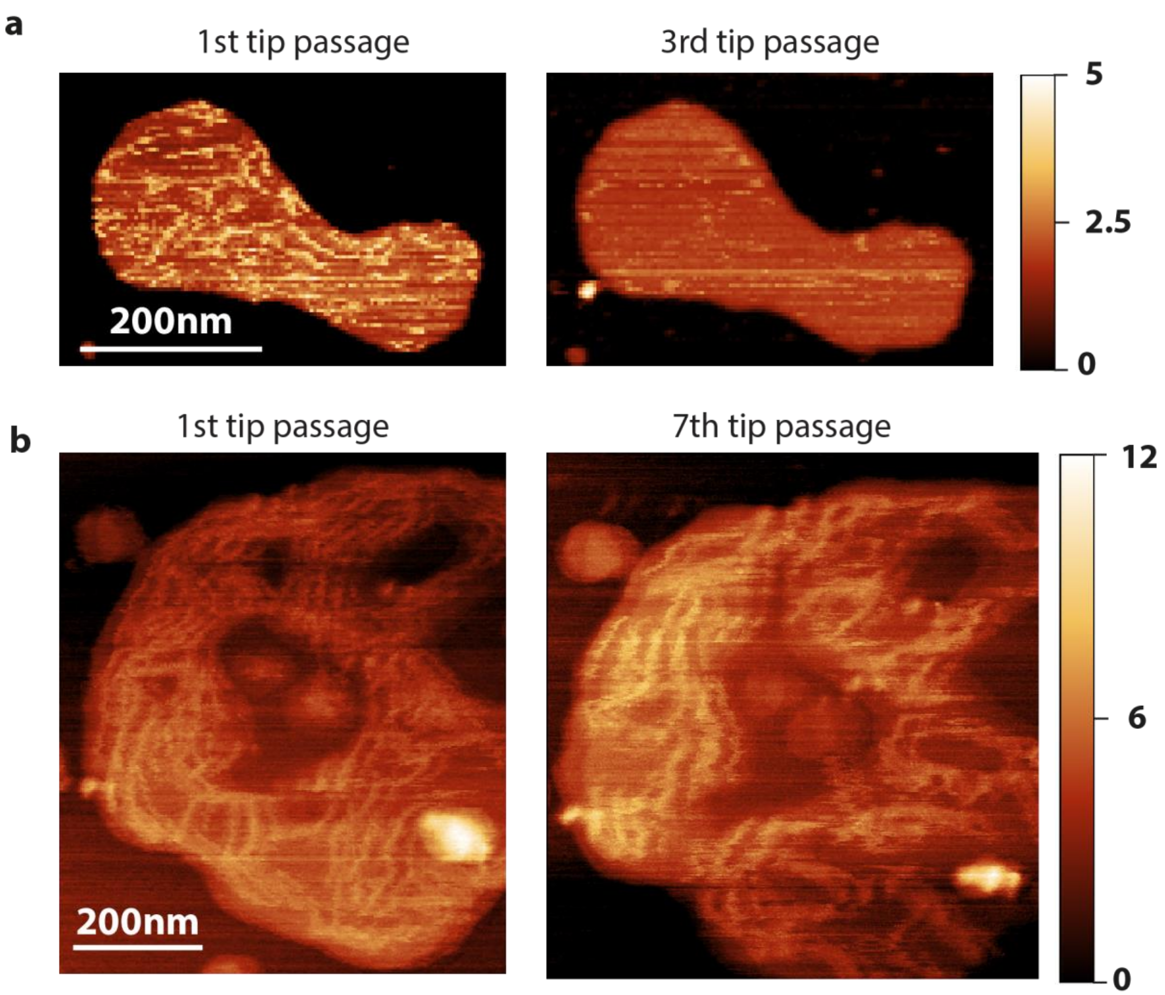


**Supplementary Figure 1: tile binding to membrane via cholesterol or aptamer anchors**. (a) Example of imaging of tiles attached to the membrane via cholesterol-oligo hybridized to the tiles. Tiles are detached by the mechanical action of the AFM tip. (b) Example of imaging of tiles attached to biotinylated lipids via biotin-aptamers that are part of the tile sequence itself.

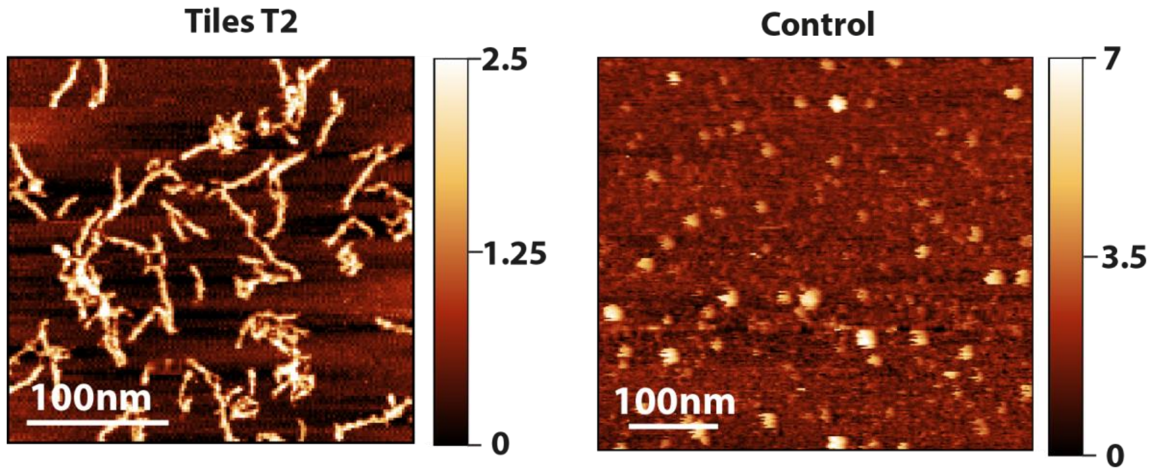


**Supplementary Figure 2: isothermal tile refolding.** 500nM of T2 tiles were incubated for 5 minutes in buffer supplemented with 12.5mM $MgCl_2$ (left panel) or 5mM EDTA (right panel) and subsequently deposited on mica for AFM imaging. Spontaneous isothermal tile refolding, resulting in formation of filaments, is visible in the presence of $MgCl_2$. In the presence of EDTA the tiles remained unfolded due to self-repulsion of the ssRNA strand, resulting in the absence of filaments.
